# Discovery of Novel Small Molecule CB2 Agonist for the Treatment of Glioblastoma Tumors

**DOI:** 10.1101/2023.12.16.572001

**Authors:** M. Kiraly, D. Okwan-Duodu, A.J. Wong, J.F. Foss, T. Giordano

## Abstract

Malignant brain tumors cause over 15,000 deaths per year in the United States. Survival for over five years is only 36%. Nearly 49% of malignant brain tumors are glioblastomas (GBM), and 30% of them have the ability to diffuse and infiltrate. Treatment frequently includes surgery, radiotherapy and chemotherapy. In case of GBM patients, combining temozolomide (TMZ) chemotherapy with radiation improved survival over radiotherapy alone (survival by 2 years: 17% vs. 11%; 5 years: 10% vs. 2%). Most primary GBM tumors from pediatric and adult patients express high levels of cannabinoid type II (CB2) receptors, and that expression correlated with tumor grade. Cannabinoids like (−)-trans-Δ^9^−tetrahydrocannabinol (Δ^9^-THC) were shown to suppress GBM tumor growth, trigger apoptosis in GBM stem cells, and slow down angiogenesis, thus cutting GBM cells off of blood supply. These data led to local administration of Δ^9^-THC in clinical trials in patients with recurrent glioblastomas, although the well-known psychotropic effects of Δ^9^ -THC and related compounds mediated via the CB1 receptors have raised some concerns among clinicians. Thus, the medicinal usage of cannabinoids has been limited. One leading strategy to avoid the side effects is administration of CB2-selective non-psychotic drugs. To create an effective solution, we designed a preclinical study to develop a novel GBM therapy, using NeuroTherapia’s lead molecule, the CB2 agonist NTRX-07. We have already demonstrated that NTRX-07 ameliorates Amyloid β production and deposition in the hippocampus, and thus restored Long-Term Potentiation – the cellular mechanism for learning and memory formation. Consequently, we showed that NTRX-07 has a highly competitive target compound profile, and that is safe in murine models, dogs and humans. NTRX-07 has entered clinical trials for the management of AD as the first orally available CB2 agonist designed to be centrally active. The phase I single ascending dose study in normal volunteers demonstrated targeted plasma levels of the drug after oral administration with no serious adverse events or clinically significant changes in safety examinations or laboratory tests. In this pilot mouse GBM survival study, we found breakthrough evidence that our compound can exert potent anti-cancer activity and significantly extend the survival of GBM animals; even without previously exposing them to radiotherapy or TMZ. Our goal is to bring NTRX-07 into the clinic as a new therapeutic for patients with GBM.

## Introduction

Malignant brain tumors cause over 15,000 deaths per year in the United States. The incidence of primary malignant brain tumors is around 7 of 100000 individuals, and this rate increases with age. Survival for over five years is only 36%. Nearly 49% of malignant brain tumors are glioblastomas (GBM), and 30% of them have the ability to diffuse and infiltrate. Treatment frequently includes surgery, radiotherapy and chemotherapy. In case of GBM patients, combining temozolomide (TMZ) chemotherapy with radiation improved survival, over radiotherapy alone (survival by 2 years: 17% vs. 11%; 5 years: 10% vs. 2%). Schaff et. al, 2023 summarizes current first-line treatments of adult-type diffuse glioma. Most patients die from progressive disease; there is therefore an urgent and important unmet need to identify novel targeted therapies.

A compelling article by Ellert-Miklaszewska et al., 2007 showed that most primary GBM tumors from pediatric and adult patients express high levels of cannabinoid type II (CB2) receptors, and that expression correlated with tumor grade. Anti-proliferative effects of cannabinoids have been reported in various cancer models *in vitro*, including malignant glioma cells (Mangal et al., 2021; Dumitru et al., 2018; De Petrocellis et al., 1998; Duntsch et al., 2005; Guzman et al., 2001; Massi et al., 2004; McAllister et al., 2005; McKallip et al., 2002; Portella et al., 2003; Ruiz et al., 1999; Sanchez et al., 1998; Sarker et al., 2003). Sanchez et al., 1998 showed that (−)-trans-Δ^9^−tetrahydrocannabinol (Δ^9^-THC), a non-specific cannabinoid receptor agonist, blocks the growth of C6 glioma cells *in vitro* and triggers apoptosis. More importantly, the antiproliferative action of cannabinoids, mediated via the CB1 and/or CB2 cannabinoid receptors, resulted in a significant regression of rat and human malignant gliomas in the Δ^9^-THC-treated animals (Duntsch et al., 2005; Galve-Roperh et al., 2000; Massi et al., 2004; Sanchez et al., 2001). These data led to clinical trials in patients with recurrent glioblastomas (Guzman et al., 2006). Unfortunately the well-known psychotropic effects of Δ^9^-THC and related compounds mediated via the CB1 receptors have raised some concerns among clinicians, thus limiting the medicinal usage of cannabinoids. One leading strategy to avoid the side effects is the administration of CB2-selective non-psychoactive drugs.

CB2 receptors (CB2Rs) were initially regarded as peripherally expressed cannabinoid receptors, because early CB2R studies demonstrated lack of detectability in the brain (Munro et al., 1993). Since then, several other basic science and clinical investigators using various molecular, imaging, electrophysiology and ultrastructural research approaches confirmed that functional CB2Rs are present in the central nervous system (CNS). A recent comprehensive review discusses these findings in detail, as well as the therapeutic roles of CB2Rs in neurodegenerative as well as psychiatric disorders, including but not limited to Alzheimer’s disease (AD), Parkinson’s disease, Huntington’s disease, anxiety, depression or schizophrenia (Kibret et al., 2022).

CB2 agonists hold the potential for breakthrough innovation in cancer therapeutics as well. As shown in Figure 1, multiple molecular pathways and mechanisms have been implicated in GBM pathophysiology, in response to cannabinoids (Dumitru et al., 2018). Briefly, there are three categories. First, the pro-apoptotic Bad protein is phosphorylated and trafficked to the mitochondria, thereby compromising the mitochondrial membrane and inducing Cytochrome C and cathepsin release, which in turn activates apoptosis-triggering caspases. This process is mediated via an increase in intracellular ceramide content that blocks the pro-survival P13K/Akt and Raf1/MEK/ERK pathways. Ceramide also activates p8/TRB3 and inhibits Akt/mTORC1, and thus enhances glioma cell autophagy (Carracedo et al., 2006; Salazar et al., 2009; Hernández-Tiedra et al., 2016, Hosami et al., 2021). Furthermore, it has been shown that cannabinoid treatment of GBM cells causes oxidative stress via the release of reactive oxygen species (ROS) that have been linked to programmed cell death induction (reviewed by Massi et al., 2010 and Hosami et al., 2021). Second, THC has proven to arrest the cell cycle of GBM cells by reducing levels of proteins promoting cell cycle progression (E2F1 and Cyclin A) and restricting proliferation via ROS production (Galanti et al., 2008; Massi et al., 2010, Hosami et al., 2021). CB2 agonist JWH133 was shown to suppress angiogenesis in malignant gliomas *in vivo*, as well as in cell culture settings (Blázquez et al., 2004). Third, cannabinoids suppress cell migration or sprouting of endothelial cells *in vitro* and *in vivo*. The molecular underpinnings for preventing proliferation, angiogenesis and invasion are the protein kinase C (PKC) and p38-MAPK pathways, combined with the downregulation of other pro-angiogenic factors such as matrix metalloproteases, as well as several growth factors (Blázquez et al., 2003, Blázquez et al., 2004, Solinas et al., 2012, Hosami et al., 2021). Lastly, researchers observed an upregulation of NKG2D ligands (MICA/B) on the GBM cell surface via STAT3 inactivation, leading to enhanced clearance of glioma cells by NK cells (Ciaglia et al., 2015).

**Fig. 1:**
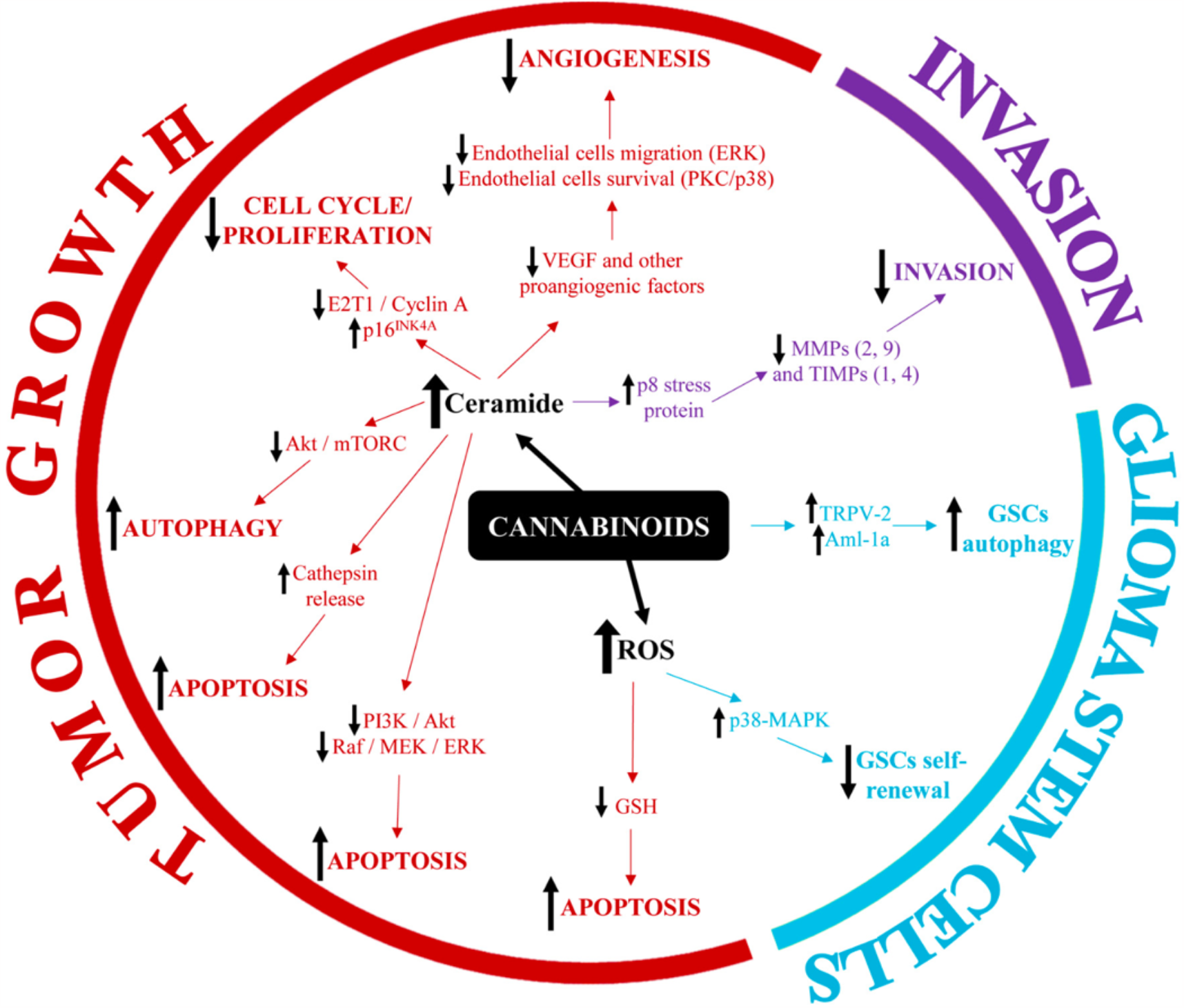
Lead mechanisms of action of cannabinoids in the treatment of GBM.

CB2Rs are G-protein coupled receptors (GPCRs), and CB2 agonists can activate them to various degrees via two separate pathways: the canonical (cAMP inhibition) and non-canonical (arrestin recruitment) cascades (Magham et al., 2021, Dopeswarkar et al., 2016). Published evidence shows that intracellular cAMP accumulation after treatment with JWH133 directly inhibits the growth of glioma cells through AMPK activation (Sánchez et al., 2001; Wang et al., 1019); whereas arrestin recruitment has been generally associated with ceramide accumulation via ERK1/2 (Magham et al., 2021, Dopeswarkar et al., 2016) that triggers cell death as discussed in the previous paragraph. Lei et al., 2020 article presents data that JWH133 even more effectively slows GBM tumor growth in combination with other compounds that prevent angiogenesis. JWH133 is an efficacious, cyclase biased agonist, nearly ineffective in arrestin-based assays (Dopeswarkar et al., 2016). NeuroTherapia’s lead compound, NTRX-07 has proven in both β-arrestin (EC50 = 23 nM) and cAMP (EC50= 19.5 nM) assays to be an efficacious unbiased agonist, thus exerting therapeutic effects via both GPCR cascades. In this study we show that that NTRX-07 is a strong candidate to develop into a standalone, potent anti-cancer therapeutic.

## Materials and Methods

The purpose of this study was to determine survival of C57BL/6 mice after the intracranial implantation of GL261 murine glioblastoma model in the presence of the lead compound NTRX-07 or vehicle. Briefly, eight-to-twelve-week-old female mice were set up with 1x10^5^ GL261 tumor cells in 0% Matrigel orthotopically intracranially (top-down). Cell injection volume was 0.01 mL/animal. Animals (n=20) were randomized into treatment groups (compound vs. vehicle control) 7 days post-implant. Body weight was measured once daily x 5 then biweekly until the end of the study. Any individual animal with a single observation of > 30% body weight loss or three consecutive measurements of >25% body weight loss was euthanized. Animals that showed full hindlimb paralysis or proptosis of eye were considered moribund. The endpoint of the experiment was moribundity or Day 45 post-implantation, whichever came first. When the endpoint was reached, the animals were euthanized. Clinical signs associated with tumor progression included impairment of hind limb function, ocular proptosis, weight loss, and neurological signs (circling, behavioral changes). Full hindlimb paralysis, ocular proptosis, moribundity, or neurological signs in conjunction with 15% weight loss were considered sufficient for euthanasia. Animals were dosed daily (qd) orally (po), and the dosing solution was prepared every day using a vehicle solution consisting of 0.5% MethocelTM A4M: 1% Soluplus® in 100 mM citrate buffer, pH 4 ± 0.2. The compound (NTRX-07-SDD) and vehicle were stored at 4°C pre-formulation and - 4°C post-formulation. NTRX-07-SDD was dosed at 1200 mg/kg, containing 300 mg/kg active compound, defined as NOAEL (No-observed-adverse-effect level) by a previous 6-months rat toxicity study. The dosing volume was adjusted accordingly for body weight. Kaplan– Meier curves for survival (OS) were evaluated and the log-rank test was used to establish the significance. Statistical analysis was performed using GraphPad Prism software and the PASW Statistics 18.0 package. P values are given as two-sided and were considered statistically significant below 0.05 (see details of the statistical analysis in Tables 1 and 2 below). All animals and cells were obtained from, and all procedures and data analyses were performed at Charles River Discovery Research Services North Carolina (IACUC ASP #: 990301). Interpretation of the results was done in collaboration with Profs. Wong and Okwan.

## Results

Mice (n=10 per group) were randomized to receive vehicle or NTRX-07. Two animals were excluded due to technical failure to inoculate tumors. - **Table 1** and **Figure 2** below summarize the results obtained from our preliminary Kaplan-Meier study. For statistical comparison of the survival curves, log-rank (Mantel-Cox) test was performed (chi square = 5.006, p =0.0253). In an alternative statistical approach, we used the Gehan-Breslow-Wilcoxon test to give more weight to deaths at earlier time points, and the difference persisted (chi square = 3.936, p =0.0473). Median survival or time to endpoint (TTE) was 18 days for the vehicle control placebo (C) and 22.5 days for the treatment group (T). Treatment with NTRX-07-SSD also increased lifespan (ILS) by 25 days, human equivalent of about Two animals were treated as euthanized for sampling for stats due to a failure to inoculate tumors. TTE = Time to endpoint. ILS = Increased life span, (T/C – C) * 100. **Fig. 2A**: The survival graph and statistics show a clear significant separation of the two (NTRX-07 and vehicle dosed) curves. As shown in **Fig 2B**, : Percent group mean body weight changes from day 1 after GL261 tumor inoculation, progressively decreasing by ∼12% in control group. In constrast, the body weight of the NTRX-07 treated animals didn’t change substantially over time (-1.6% on Day 17), consistent with efficacy and tolerance of the drug. whereas that of the control group progressively decreased (-12.0% on Day17).

| Group | n | Treatment Regimen |  |  |  | Median TTE | ILS | Statistical Significance |
| --- | --- | --- | --- | --- | --- | --- | --- | --- |
|  |  | Agent | mg/kg | Route | Schedule |  |  |  |
| 1 | 8 | vehicle | --- | po | qd to end | 18.0 | --- | --- |
| 2 | 10 | NTRX-07-SDD | 300 | po | qd to end | 22.5 | 25 | * |

**Fig. 2:**
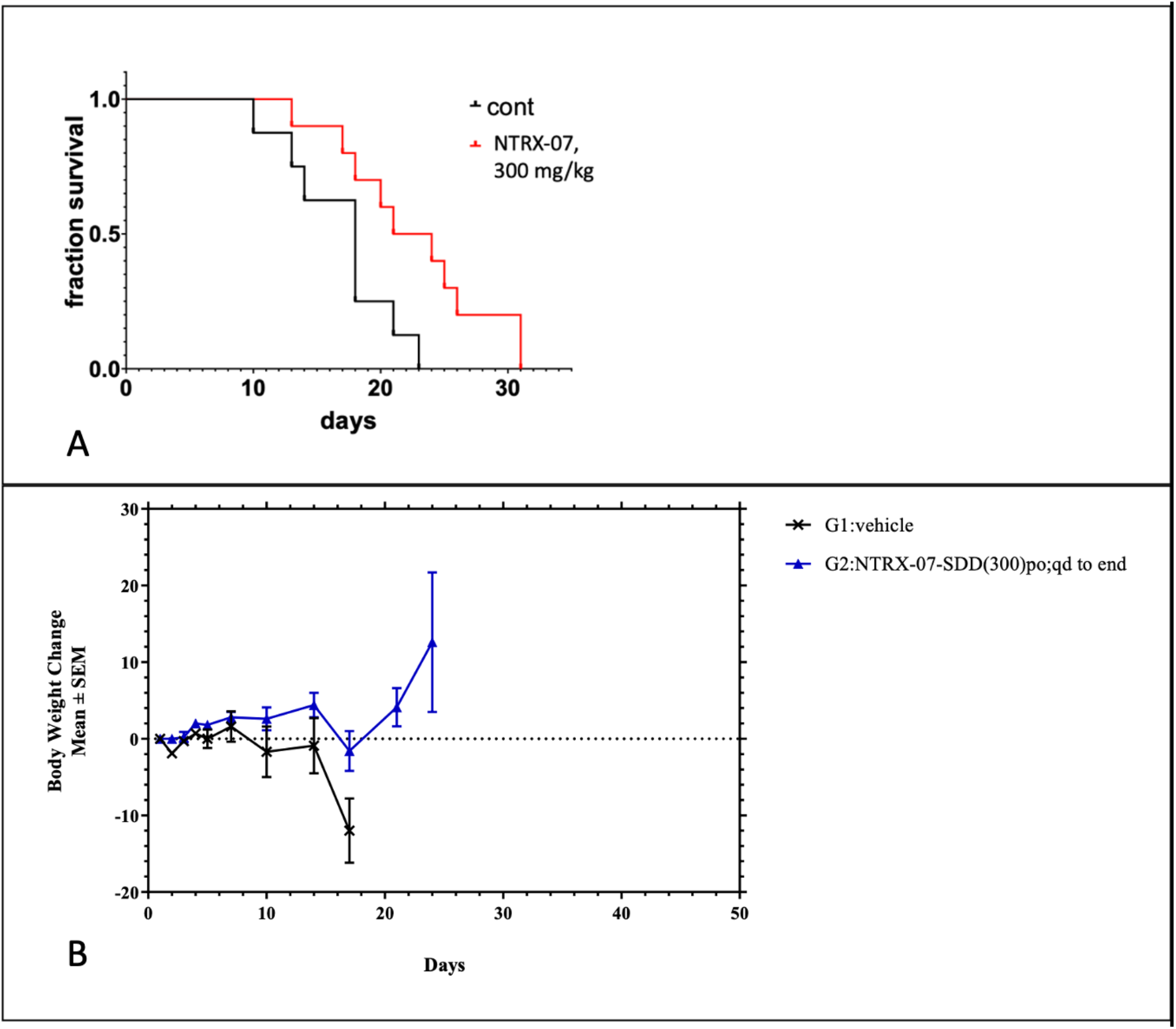
Results of our preliminary Kaplan-Meier survival study, vehicle vs. 300 mg/kg NTRX-07 treatment.

## Discussion

Our results show that 300 mg/kg of NTRX-07 is well tolerated in mice and leads to improved survival over control mice challenged with GL26 tumors. These key findings are breakthrough evidence that an orally available CB2 agonist can exert potent anti-cancer activity with the promise of prolonging life, even without previously exposing the subjects to radiotherapy. Our goal is to develop and optimize a novel therapeutic strategy using our CB2 agonist NTRX-07, that is suitable for advancement towards further (pre)clinical testing. This will be achieved by establishing the most effective dose in combination with radiotherapy and TMZ; by confirming robustness across various implanted human GBM cell xenografts; and by understanding the treatment’s strengths and weaknesses via unbiased single-cell RNAseq analysis of GBM tissue samples resected from NTRX-07/ vehicle treated animals. We anticipate that radiotherapy and TMZ, combined with NTRX-07 treatment will further increase the survival time of GBM animals, to highly statistically significant levels, both in a SCID mouse/ human patient xenograft model and the C57BL/6 /GL261 model systems. We have initiated studies at the highest dose in rodents and future studies will explore dose exposure responsiveness of the tumors. Recent reviews (Hohmann et al., 2021; Ellert-Miklaszewska et al., 2021) suggest that human GBM cells respond even more sensitively to CB2 treatment than the GL261 mouse cell line. We expect to see a maintenance of body weight, a major reduction in tumor size, decreased viability of cancer cells, inhibited angiogenesis, fewer migrating GBM stem cells and larger populations of infiltrating microglia. We believe that PK values will correlate with the dose levels, similar to what we have seen in previous rodent toxicology experiments. It is possible however that we will detect higher than usual NTRX-07 levels in brain lysates of treated GBM animals, due to their compromised blood brain barrier. However, we cannot predict how the survival curves will change, as we combine our compound with radiotherapy and TMZ. New survival curves will be needed for all the planned subsequent studies. Overall, our findings provide evidence that a CB2-selective non-psychoactive drug may improve outcomes in GBM.

